# Sex-Specific Cytokine, Chemokine, and Growth Factor Signatures in T1D Patients and Progressors

**DOI:** 10.1101/2024.09.05.611513

**Authors:** Khyati Girdhar, Keiichiro Mine, Jeffrey M. DaCosta, Mark A. Atkinson, Johnny Ludvigsson, Emrah Altindis

**Affiliations:** Boston College Biology Department, Chestnut Hill, Massachusetts, USA; Department of Pathology, Immunology and Laboratory Medicine, Diabetes Institute, College of Medicine, University of Florida, Gainesville, Florida, USA; Department of Pediatrics, Diabetes Institute, College of Medicine, University of Florida, Gainesville, Florida, USA; Crown Princess Victoria Children’s Hospital, Division of Pediatrics, Department of Biomedical and Clinical Sciences, Linköping University, Linköping, SE, Sweden

**Keywords:** Type 1 diabetes, sex, cytokine, chemokine, growth factor, autoimmunity

## Abstract

While studies have reported altered levels of cytokines in type 1 diabetes (T1D) patients, the results are inconsistent, likely because of variable factors. This study tests the hypothesis that there are sex-based differences in cytokine levels in T1D, prior to and after disease onset. We analyzed 48 blood cytokine, chemokine, and growth factor levels using a multiplex assay. We found only two cytokines, M-CSF and IL-6, with significant differences between T1D patients (n=25) versus controls overall (n=25). However, we identified notable alterations when comparing sex-age-matched controls and T1D samples. Inflammatory cytokines (TNF-α, IL-6, IL-1a), Th2 cytokines (IL-4, IL-13), and chemokines (MIP-1α, RANTES, MIP-3) were lower in female T1D patients compared to female controls, but not in males. IL-22 was lower in female T1D patients compared to female controls, while it was higher in male T1D patients compared to male controls. In contrast, growth factors (EGF, PDGF-AB/BB) were higher in male T1D patients compared to male controls. In T1D progressors (children who developed the disease years after the sample collection, n=16-21), GROa was lower compared to controls in both sexes. Our findings underscore the importance of understanding sex-specific differences in T1D pathogenesis and their implications for developing personalized treatments.

## Introduction

### T1D is an autoimmune disease characterized by pancreatic β-cell destruction

Unlike most of the other autoimmune diseases where the majority of patients are females ^1^, there is approximately 1.5 times higher incidence of T1D in males than in females ^2^. Sex is also associated with the risk of T1D complications. Males with T1D have a higher risk of advanced retinopathy and diabetic kidney disease ^3,4^, whereas females with T1D have a higher risk of cardiovascular events ^5^. Additionally, compared to male patients, females with T1D are more prone to develop multiple autoimmune diseases, including hypothyroidism, celiac disease, and Addison’s disease ^6^.

Sex is an important factor in regulating immune responses. Circulating CD4/CD8 T cell ratios are higher in females versus males in the general population ^1^. As for cytokine secretion, peripheral blood mononuclear cells (PBMCs) from healthy females secrete higher levels of interferon (IFN)-α in response to toll-like receptor (TLR) 7 ligands than those from healthy males ^7^; whereas PBMCs from healthy males produced higher interleukin (IL)-10 levels in response to TLR9 ligand or herpes simplex virus type 1 infection than those from healthy females ^8^. It is suggested that sex chromosomes drive these differences in cytokine secretion ^7–9^. These sex-specific immune responses and the association of sex in T1D suggest sex may be a crucial factor in the disease pathogenesis.

Several studies have investigated the role and level of cytokines - key modulators of the immune responses and immunotherapeutic targets ^10^ - using blood samples (serum or plasma) from patients with T1D. Studies on T1D patients report varying results. For example, Fatima et al. reported IL-1β and IL-17A ^11^, whereas Svensson et al. reported transforming growth factor (TGF)-β and IL-18 were elevated in T1D patients ^12^. Conversely, some studies have reported more than ten cytokines, including proinflammatory and Th17 cytokines, as elevated in T1D patients ^13,14^. These differences in reported cytokine levels may be due to the potential influence of factors including age, disease duration, and race/ethnicity ^13^. In addition, sex was suggested to be associated with cytokine levels in T1D patients, as males had higher TNF-α levels than females ^12,13^. However, few studies have focused on sex differences among these patients.

In this small, exploratory study, we tested the hypothesis that male and female T1D patients and progressors have distinct cytokine signatures compared to healthy age- and sex-matched controls. To this end, we evaluated blood samples from two different human cohorts: 1) T1D patients and matched controls from the University of Florida Diabetes Institute (UFDI) Study Bank ^15,16^ and 2) five-year-old T1D progressors (diagnosed years after sample collection) and age-sex-matched controls from All Babies in Southeast Sweden (ABIS) cohort ^17,18^.

## Results

### T1D patients have sex-dependent serum cytokine, chemokine, and growth factor profiles

Using a multiplex assay, the levels of 48 cytokines, chemokines, and growth factors were measured in serum samples obtained from 25 T1D patients (12 males, 13 females) and 25 age-sex-matched healthy controls (**Fig. 1A, Table 1, and Supplementary Table 1,** UFDI Study Bank cohort). We identified only two cytokines, macrophage colony-stimulating factor (M-CSF) (2.0-hold) and IL-6 (1.9-hold), which showed a significant decrease in T1D versus controls overall (**Fig. 1B and 1C**). However, when we analyzed these biomarker levels according to sex, we identified significant sex-specific differences. Inflammatory cytokines, including tumor necrosis factor (TNF)-α (2.1-fold), IL-6 (2.4-fold), and IL-1a (4.1-fold), were lower in female T1D patients compared to female controls, but not in males (**Fig. 1C, 1D**, and **1E**). Similarly, amongst females, the Th2 cytokines IL-4 (2.3-fold) and IL-13 (1.8-fold) were lower in T1D versus controls, but there were no differences in males (**Fig. 1F** and **1G**). Notably, IL-22, a cytokine involved in mucosal immunity ^19^, was 2.4-fold lower in female T1D patients compared to female controls, while it was 1.9-hold higher in male T1D patients compared to male controls (**Fig. 1H**). Chemokines including macrophage inflammatory protein (MIP)-1α (also known as CCL3, 2.0-fold), regulated upon activation, normal T cell expressed and secreted (RANTES) (also known as CCL5, 3.9-fold), and MIP-3 (CCL7, 2.6-fold) were lower in female T1D patients compared to female controls, while there were no differences in males (**Fig. 1I, 1J**, and **1K**). Interestingly, all differences observed in growth factor levels were specific to the male participants. Epidermal growth factor (EGF) (1.6-fold) and platelet-derived growth factor-AB/BB (PDGF-AB/BB) (1.1-fold) were higher in male T1D patients compared to male controls (**Fig. 1L** and **1M**).

**Figure 1:**
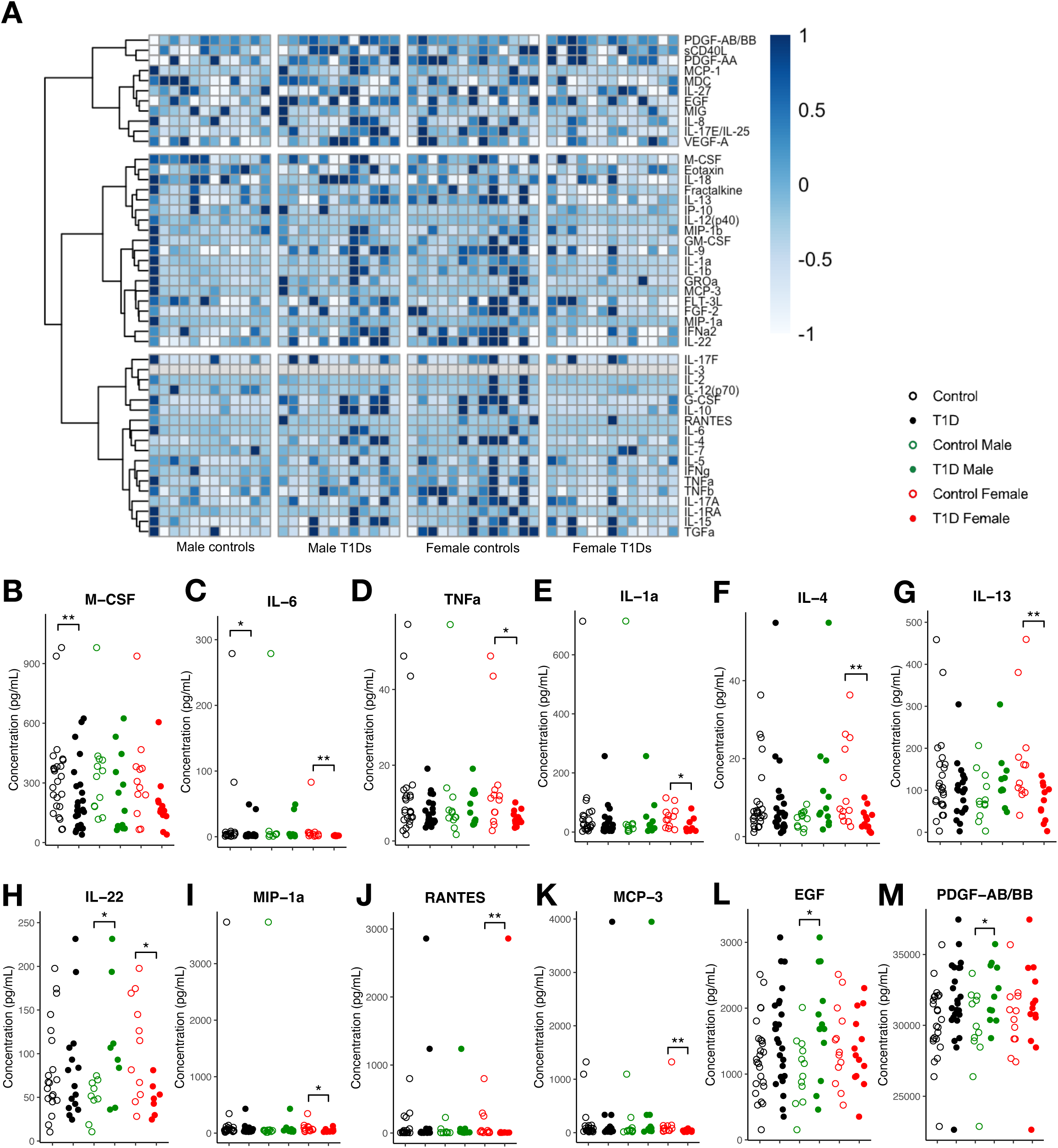
The sex-based differences of cytokine profiles in T1D patients. (A) Heatmap showing all 48 cytokine levels. (B-M) Cytokine levels in the serum of T1D patients in the UoF cohort (n=25; male, n=12; female, n=13) and healthy controls (n=25; male, n=12; female, n=13). Statistical significance was evaluated using Mann-Whitney *U*-tests with a false discovery rate correction for multiple comparisons (\**P*<0.05 and *Q*<0.1; \*\**P*<0.05 and *Q*<0.05).

**Table 1.** Characteristics of T1D patients and healthy controls.

| Characteristics | Type 1 diabetes | Healthy controls |
| --- | --- | --- |
| Participant number | 25 (12 males, 13 females) | 25 (12 males, 13 females) |
| Age - yr (range) | 18.1 ± 4.1 (10-25) | 18.1 ± 4.1 (10-25) |
| Male | 17.5 ± 3.7 | 17.5 ± 3.7 |
| Female | 18.6 ± 4.4 | 18.6 ± 4.4 |
| Age at diabetes onset (range) | 10.5 ± 5.4 (1.8-19.2) | - |
| Male | 9.9 ± 4.8 | - |
| Female | 11.0 ± 5.9 | - |
| Disease duration - yr (range) | 8.1 ± 6.5 (1.0-22.6) | - |
| Male | 8.1 ± 5.6 | - |
| Female | 8.1 ± 7.2 | - |
| BMI (kg/m<sup>2</sup>) | 23.3 ± 3.6 | 25.9 ± 6.4 |
| Male | 23.1 ± 3.8 | 26.0 ± 5.6 |
| Female | 23.4 ± 3.4 | 25.8 ± 7.0 |

Values are means ± standard deviation.

In multiple regression analysis, we identified very few samples. Only PDGF-AA/BB levels were correlated with body mass index (BMI) (rho, 0.30; **Supplementary Fig 1A**), and M-CSF levels decreased with age (rho, −0.29; **Supplementary Fig 1B**). Although, we had a small sample size, PDGF-AA/BB was associated with race with higher levels detected in participants who reported their race as Asian as compared with Black/African American (**Supplementary Fig 1C**).

### Growth-regulated alpha (GROα) is decreased in T1D progressors years before diagnosis

Our findings showed that T1D patients have sex-based cytokine signatures. To determine if these differences exist before overt T1D onset, we utilized plasma samples obtained from 5-year-old T1D progressors (ABIS cohort) (**Fig. 2A**). Using these pre-adolescent samples (16 T1D progressors and 21 healthy controls), we were able to minimize the impact of established disease, disease-related therapies, and sex hormone activity (**Table 2 and Supplementary Table 2**).

**Figure 2:**
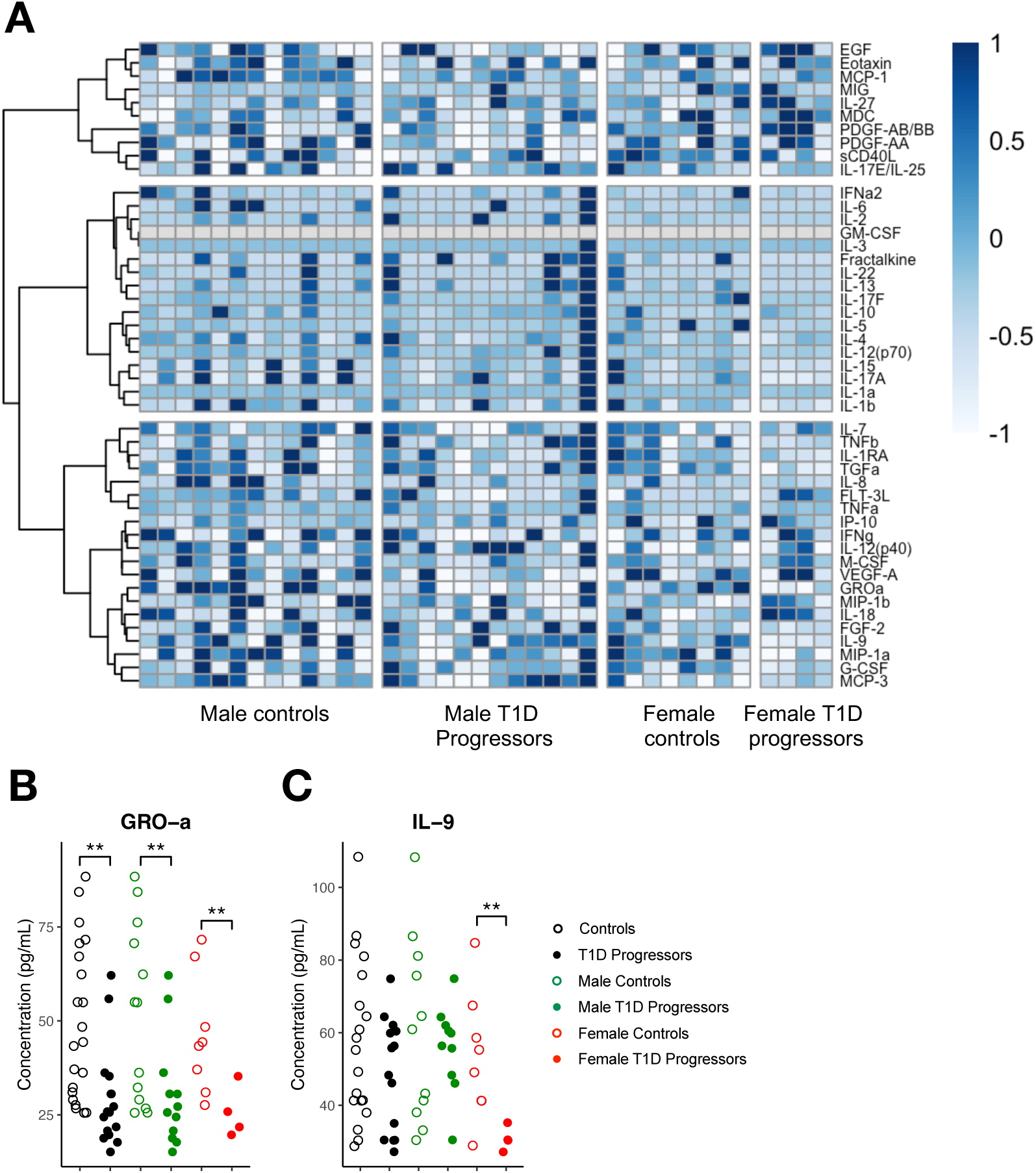
The sex-based differences of cytokine profiles in T1D progressors. (A) Heatmap showing all 48 cytokine levels. (B, C) Cytokine levels in the plasma of T1D progressors in the ABIS cohort (n=16; male, n=12; female, n=4) and healthy controls (n=21; male, n=13; female, n=8). Statistical significance was evaluated using Mann-Whitney *U*-tests with a false discovery rate correction for multiple comparisons (\**P*<0.05 and *Q*<0.1; \*\**P*<0.05 and *Q*<0.05).

**Table 2.** Characteristics of T1D progressors and healthy control.

| Characteristics | T1D progressors | Healthy controls |
| --- | --- | --- |
| Participant number | 16 (12 males, 4 females) | 21 (13 males, 8 females) |
| Age of sample collection | 5 ± 0 | 5 ± 0 |
| Age of diagnosis - yr (range) | 14.6 ± 4.0 (6.8-21.7) | - |
| Male | 14.9 ± 4.3 | - |
| Female | 13.6 ± 2.0 | - |

Values are means ± standard deviation.

GROα (also known as CXCL1) was 1.7-fold lower in all T1D progressors compared to all healthy controls (**Fig. 2B**). This decrease was not sex-specific and observed in both male (2.1-fold) and female (1.8-fold) T1D progressors (**Fig. 2B**). On the other hand, while our sample size was quite low, IL-9 was 1.7-fold lower in female progressors only (**Fig. 2C**). Interestingly, these alterations were specific to the T1D progressors, and they were not observed for T1D patients in the UFDI Study Bank cohort.

## Discussion

Similar to other autoimmune diseases ^20^, cytokines are potential immunotherapeutic targets for T1D because of their central role in orchestrating the immune response ^10^. In the preclinical animal model of T1D, non-obese diabetic (NOD) mice, treatment of cytokines, including IL-4, IL-13, IL-17E/IL-25, or IL-33, suppressed the diabetes onset [10, 19]. On the other hand, deficiency of IL-10, an anti-inflammatory cytokine, accelerated diabetes onset as expected ^21^. In humans, treatment with inhibitor-targeted Janus kinase (JAK) 1 and 2, non-receptor tyrosine kinases involved in signaling downstream of several cytokines, was reported to preserve β-cell function in new-onset T1D patients ^22^. These observations suggest that cytokines play an important role in the disease pathogenesis.

Sex is a crucial biological variable in immunological research, affecting both innate and adaptive immune responses ^1^. For example, Plasmacytoid dendritic cells (pDCs) from females secrete higher IFN-α levels than those from males when stimulated with TLR-7 ligands ^23^. Cytokine production levels also vary between males and females in the context of infection ^24^. However, there has been limited data focusing on sex differences in cytokine levels in T1D studies. Earlier studies reported higher *IFN-*γ and *IL4* mRNA levels in PBMCs from males with T1D compared to male controls, with no difference in females ^25^. In newly diagnosed T1D cases, plasma TNF-α levels were higher in males than females while females had higher levels of IL-1Ra ^13,26^. This evidence highlights the importance of sex as a variable in T1D research.

Th2 cytokines such as IL-4 and IL-10 play a role in suppressing immune responses to maintain homeostasis ^10^. Interestingly, in our study, Th2 cytokines, IL-4 and IL-13, were lower in females with T1D than in female controls while there was a trend of increase in male patients. Previous reports demonstrated IL-4 treatment suppressed β-cell apoptosis via activation of the PI3K pathway, while IL-13 impaired IFN-γ secretion from splenocytes, leading to reduced diabetes onset in female NOD mice ^27–29^. These observations suggest that decreased Th2 responses in female patients may contribute to T1D pathogenesis. Therapies increasing Th2 cytokines could be beneficial in slowing β-cell destruction and/or diabetes related complications.

Th1 cytokines such as IL-2, IL-12, and TNF-α have a role in exacerbating autoimmunity leading to T1D by promoting the development of islet autoreactive Th1 cells and cytotoxic CD8^+^ T cells ^10,30^. In contrast to previous meta-analyses that reported higher levels of TNF-α in Type 1 diabetes (T1D) patients compared to healthy controls across both sexes ^14^, our findings indicate that TNF-α levels were lower only in females with T1D compared to female controls. These results suggest that a larger sex-specific study is needed to draw such conclusions. Finally, all biomarkers identified in this study that were significantly altered in female T1D patients were lower than in controls, which may indicate an overall systemic impairment of cytokine production specific to female T1D patients.

Some of the most interesting data obtained in our study was related to IL-22, which plays a key role in intestinal barrier protection by inducing mucus, antimicrobial protein, and complement production from intestinal epithelial cells, directly influencing the gut microbiota composition ^19,31^. Indeed, IL-22-deficient mice showed less intestinal microbiota diversity and weakened barrier function, resulting in increased susceptibility to colitis and infection ^19^. We and others have shown that the gut microbiome composition and diversity are associated with autoimmune diseases, including T1D ^32–35^, and affected by sex ^36–38^. Interestingly, *Phocaeicola vulgatus* (formerly *Bacteroides vulgatus*), a gut commensal known to reduce IL-22 levels in the intestine and serum of colonized female mice ^39^, has been reported to be increased in children at high genetic risk for T1D before seroconversion ^40^. Although IL-22 treatment itself was not effective in preventing T1D in female NOD mice ^41^, it has been reported that IL-22 signaling has a role in ameliorating T1D in NOD mice treated with fermentable dietary fiber ^42^. In our study, IL-22 showed a similar trend to that of Th2 cytokines. Specifically, IL-22 was 1.9-fold lower in females with T1D compared to female controls, whereas IL-22 was 2.4-fold higher in males with T1D compared to male controls, suggesting sex-specific IL-22 contribution to the pathogenesis of the disease.

Growth factors, including EGF and PDGF are known to be associated with the development of diabetic retinopathy ^43^. In addition, PDGF was reported to be increased in T1D patients ^44^. We observed increased EGF (1.6-fold) and PDGF-AB/BB (1.1-fold) in males with T1D versus male controls, but not in females. This observation may be related to the fact that advanced retinopathy is more common in male T1D patients ^3^.

In the ABIS cohort, GROα (CXCL1) was significantly lower in T1D progressors compared to controls, irrespective of sex. GROα is a chemokine that plays a role in inflammation by inducing angiogenesis and neutrophil recruitment ^45^. In contrast to our results in T1D progressors, elevated serum GROα was previously reported in T1D patients ^46^. In addition, female T1D progressors had lower IL-9 than female controls. While these findings should be replicated with larger cohorts to obtain conclusive results, our results indicate that low GROα may serve as a predictive marker for both sexes, and low IL-9 may specifically serve as a predictive marker for female T1D progressors.

Taken together, our study underscores the critical role of sex as a significant variable in T1D autoimmunity, with discernible differences in cytokine levels between T1D and control participants observed only when stratified by sex. However, the small sample size is a notable limitation that may impact the generalizability of our findings. To address this, follow-up studies with large sample sizes are needed. In addition, further research is required to elucidate the molecular mechanisms linking cytokines to sex-specific outcomes in T1D. Determining sex-specific cytokine profiles will advance our comprehension of T1D pathophysiology and could lead to the development of targeted, sex-specific therapeutic interventions for diagnosis, prevention, and management.

## Methods

### Human samples

Blood was obtained from subjects following the provision of written informed consent (and assent in the case of minors) under institutional review board–approved protocols (University of Florida IRB# 201400703) and the Declaration of Helsinki. The ABIS study was approved by the Research Ethical Committees of the Faculty of Health Science at Linköping University, Sweden (Dnr Linköping 287-96, Linköping 03-092, and Linköping 2018/380-32), and the Medical Faculty of Lund University, Sweden (Dnr Lund 83-97). The samples were deidentified before use in this study.

Fifty cross-sectional serum samples were obtained from the UFDI Study Bank ^15,16^: 12 male and 13 female participants with T1D (disease duration mean ± standard deviation [SD]: male 8.1 ± 4.1 years, female 8.1 ± 4.1 years) as well as 12 male and 13 female age-sex-matched healthy controls. Thirty-seven plasma samples were obtained from the ABIS cohort ^17,18^: 16 T1D progressors (12 males, 4 females) and 21 healthy controls (13 males, 8 females), all collected at age 5. The patient demographics are outlined in **Supplementary Tables 1 and 2**.

### Cytokine measurement

Cytokine levels were measured using the Milliplex Human Cytokine/Chemokine/Growth Factor Panel A (Millipore). Briefly, samples were thawed at 4°C prior to assay and kept on ice throughout the assay procedures. The manufacturer’s protocols were followed, and the general steps are outlined. All kit components were brought to room temperature. Reagents, including the wash buffer, beads, and standards, were prepared according to the kit instructions. The assay plate (96-well) was pre-wetted with 200 μL of wash buffer, and the solution was decanted. Next, 25 μL of each standard or control solution was dispensed into their respective wells. The sample wells received 25μL of assay buffer. For the background, standards, and control wells, 25 μL of serum matrix solution was added. Next, 25 μL of serum samples, diluted 1:3 for ABIS cohort in assay buffer, were placed into the corresponding wells. Finally, 25 μL of premixed beads were added to every well and incubated at 4°C overnight (16-18 hours) on a plate shaker (500rpm). After primary incubation, the plate was washed twice. A 25 μL detection antibody cocktail was then added to all wells. The plate was covered and incubated at room temperature for one hour on a plate shaker. After the one-hour incubation, a 25 μL streptavidin-phycoerythrin fluorescent reporter was added to all wells, and the plate was covered and incubated at room temperature for 30 minutes on a plate shaker. The plate was then washed twice. Beads were resuspended in 150 μL sheath fluid, placed on a shaker for 5 minutes, and then read on Bio-Plex® 200 following the manufacturer’s specifications and using Bio-Plex Manager software version 6.2. Multiplex protein analysis was performed by the Multiplex Core Facility at The Forsyth Institute (Cambridge, MA). In the UF cohort, 48 cytokines were analyzed. However, in the ABIS cohort, 47 cytokines were analyzed because we were unable to measure RANTES due to limitations in sample volume.

### Data analysis

Statistical analyses were performed using R v4 and nonparametric tests due to considerable skew across groups. For each cytokine, Mann-Whitney *U*-tests were used for separating pairwise comparisons (all samples, only males, only females) of T1D and control patients, with a false discovery rate correction (conversion of *P*-values to *Q*-values) for running three tests. Statistical significance for these tests were determined using two thresholds: a *P*-value<0.05 and a *Q*-value<0.1 or both *P*- and Q-values below 0.05 (* and **, respectively). For testing of other factors that may contribute to cytokine levels, numerical variables (Age and BMI) were analyzed with Spearman rank correlation tests and categorical variables were analyzed with Mann-Whitney *U*-tests (Ethnicity) or Kruskal-Wallis tests (Race). These tests were considered significant at an alpha level of 0.05.

## Acknowledgements

We thank the Multiplex Core at The Forsyth Institute (Cambridge, MA), which provided all multiplex protein analysis services. We thank Amanda L. Posgai for her assistance with editing. Finally, we extend our gratitude to all the individuals who provided samples for this study.

## Author contributions

Conceptualization, E.A.; Experiments, K.G.; Writing - original draft preparation, K.M.; Writing - review and editing, K.G., K.M., J.M.D., M.A.A., J.L., J.D., and E.A.; Supervision, E.A.; Project administration, E.A.; Statistical analysis, J.M.D.; Resources, E.A., M.A.A., J.L.; Funding acquisition, E.A., M.A.A., J.L.. All authors have critically reviewed and edited the manuscript and finally agreed to the published version of the manuscript.

## Competing Interests

The authors declare no conflict of interest. The funders had no role in the writing of this manuscript.

## Data availability statement

The all data analyzed in this study are in Supplementary tables.

## Ethics declarations

The UFDI Study was approved by the University of Florida (IRB# 201400703) and the Declaration of Helsinki. The ABIS study was approved by the Research Ethical Committees of the Faculty of Health Science at Linköping University, Sweden (Dnr Linköping 287-96, Linköping 03-092, and Linköping 2018/380-32), and the Medical Faculty of Lund University, Sweden (Dnr Lund 83-97).

## Funding

This work was supported by Beatson Foundation grant to EA, Manpei Suzuki Diabetes Foundation to KM, Diabetes Research Connection to KG, and NIH P01 AI042288 to MAA. The ABIS-study has been supported by the Swedish Research Council (K2005-72X-11242-11A and K2008-69X-20826-01-4) and the Swedish Child Diabetes Foundation (Barndiabetesfonden), JDRF Wallenberg Foundation (K 98-99D-12813-01A), Medical Research Council of Southeast Sweden (FORSS), Swedish Council for Working Life and Social Research (FAS2004-1775), Östgöta Brandstodsbolag, and ALF Region Östergötland and Linköping university, and Joanna Cocozza Foundation.

## Notes

### Competing Interest Statement

The authors have declared no competing interest.

