## Supplementary figures and images for "Sex-Specific Cytokine, Chemokine, and Growth Factor Signatures in T1D Patients and Progressors"

### Supplemental figures

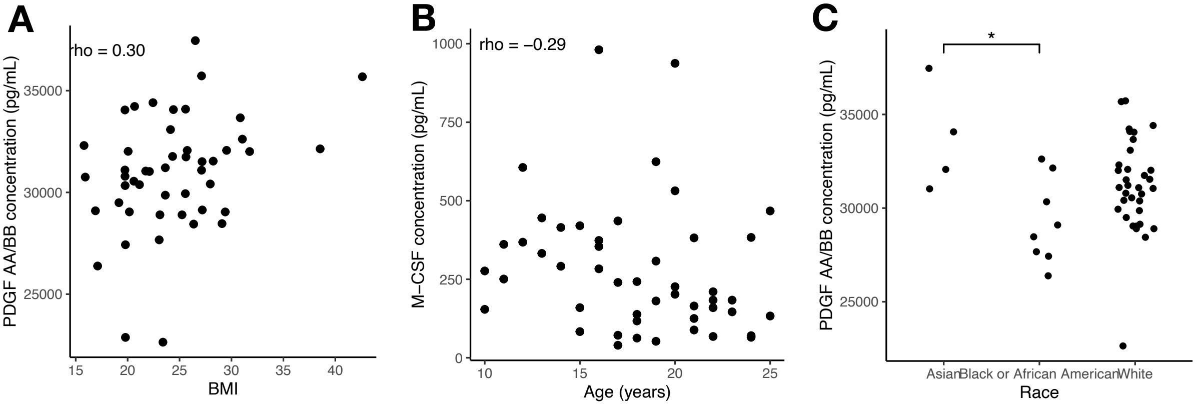


**Supplementary Figure 1: multiple regression analysis in T1D patients.**
